# Anterior cingulate cortex activation of claustrum projection neuron subtypes is enhanced by alcohol

**DOI:** 10.64898/2026.01.19.700421

**Authors:** Andreas B. Wulff, Samuel H. Sheats, Eliza A. Douglass, Brian N. Mathur

## Abstract

Cognitive impairment is a major component of Alcohol Use Disorder. Optimal cognitive performance requires anterior cingulate input activation of the claustrum, a subcortical nucleus that orchestrates cortical activity. Yet the impact of chronic alcohol exposure on the ability for the anterior cingulate cortex to drive activity of claustrum projection neuron subtypes is unknown. In adult male and female mice, we found that the majority of non-burst firing Type 1 claustrum projection neurons did not express the vesicular glutamate transporter 2 (VGLUT2), while the majority of burst firing Type 2 projection neurons were VGLUT2-expressing. Following chronic intermittent vaporized ethanol exposure (CIE), we found that both Type 1 and VGLUT2-non-expressing neurons exhibited increased responsivity to anterior cingulate cortex input activation that was mediated by increased postsynaptic membrane excitability. In contrast, Type 2 and VGLUT2-expressing projection neurons exhibited increased responsivity to anterior cingulate cortex input due to strengthened pre- and post-synaptic transmission mechanisms. Altogether, we uncovered a hyper-excitatory drive of the claustrum by the anterior cingulate cortex following chronic alcohol exposure. The data provide a foundational resource for the complex effects of chronic alcohol exposure on the claustrum, a critical cognitive control nucleus.

## Introduction

Alcohol is the most common substance of misuse contributing to more than half of all substance use disorders in the United States [1]. Alcohol use disorder (AUD) is defined by a compulsive-like consumption of alcohol driven, in part, by compromised cognitive control [2]. The claustrum is a subcortical telencephalic nucleus that, by virtue of its extensive projections back to the cortical mantle, is implicated in cognition [3–5]. Indeed, the evidence for involvement of the claustrum in attention [6], decision-making [7], and cognitive performance [3,8] in awake animals as well as memory consolidation during sleep [9,10] suggests a broad role for this structure in cognitive control [5]. Implicating the claustrum in cognitive dysfunction in substance use disorders, claustrum activity is impacted by cocaine [11–13], methamphetamine [14], and opioids [15,16], and is causally associated with control of drug seeking [11–17]. However, the impact of alcohol on claustrum synaptic transmission and excitability remains unknown.

Cognitive control impairment in AUD is characterized by structural and functional changes in the prefrontal cortices, including the anterior cingulate cortex (ACC) [2,18,19]. In rodents, the dorsomedial prefrontal cortex, including the ACC, governs alcohol intake [20–22] and these regions are sensitive to the effects of chronic alcohol exposure [23–28], as are their downstream projection targets [3,29,30]. The claustrum receives among the strongest cortical inputs from the ACC, which supports performance in cognitively demanding tasks [3,4]. The sensitivity of the claustrum to drugs of abuse and the established negative impact of chronic alcohol use on cognition suggest that ACC activation of the claustrum is sensitive to alcohol.

Discrete ethanol-induced changes to neuronal subtypes in several brain regions give rise to distinct behavioral and circuit consequences [31–34]. In the claustrum, subtypes of neurons exist based on molecular [35,36] and physiological properties [37,38] and exhibit different projection [35–37] and activity profiles [4,37,38] that are positioned to guide distinct cognitive functions [8,39]. Subtypes of projection neurons have been well-characterized in the claustrum.

Claustrum projection neurons are defined physiologically as non-burst firing “Type 1” and burst firing “Type 2” neurons and can be molecularly profiled as vesicular glutamate transporter 2 (VGLUT2)-expressing (VGLUT2+) or VGLUT2-non-expressing (VGLUT2-) [4,35–37]. Here, we determined that Type 2 neurons largely express VGLUT2, while the majority of Type 1 neurons lack VGLUT2 expression. Examining ACC input to the claustrum, we discover that chronic intermittent vaporized alcohol exposure increases ACC excitatory drive of claustrum circuitry in mice through different mechanisms based on neuronal subtype. These data lay the groundwork for understanding the complex effects of alcohol on this cognitive control nucleus.

## Materials and Methods

### Animals

All experiments were performed in accordance with NIH guidelines and were approved by the Institutional Animal Care and Use Committee of the University of Maryland Baltimore. Transgenic male and female VGLUT2-cre mice (Jax#028863) were bred on a C57BL/6J background and housed under a 12-hour light/dark cycle (lights on at 0700 hours, off at 1900 hours) with ad libitum access to food and water. All patch-clamp experiments were performed during the light cycle. Mice were 12-18 weeks of age at the time of electrophysiology experiments, which were performed during the light cycle.

### Viral Surgeries

All surgeries were performed under isoflurane anesthesia in a Kopf Stereotaxic frame. Viruses were injected using a Hamilton syringe at a rate of 10-30nL/min. All mice received bilateral injections of 100-250nL AAV5-hSyn-DIO-mCherry (Addgene #50459_AAV5) to the claustrum (AP +0.9mm, ML ±2.75mm, DV -4.0mm). A subset of mice simultaneously received bilateral injections of 250nL AAV5-hSyn-hChR2(H134R)-EYFP (Addgene #26973_AAV5) into the ACC (AP +1.0mm, ML ±0.3mm, DV -1.1mm). In total 5-6 weeks were given for viral expression before mice were euthanized for patch-clamp electrophysiology.

### Chronic Intermittent Ethanol (CIE) vapor exposure

Following 3-5 days of post-surgery recovery, mice were randomly assigned to either air or ethanol vapor exposure. Mice were exposed to either air or ethanol vapor for 16 hours/day, 4 days/week for a total of 5 weeks. Ethanol vapor exposure was performed as per our previously published protocol [40]. During week 4-5 of vapor exposure mice received pyrazole (68.1 mg/kg, I.P.) prior to start of exposure. Following the last ethanol exposure mice went through forced abstinence for 64-160 hours before being euthanized for electrophysiological recordings.

### Acute slice preparation

250μm thick coronal slices were prepared in ice-cold sucrose-modified artificial cerebrospinal fluid (in mM: 194 sucrose, 30 NaCl, 4.5 KCl, 1 MgCl_2_, 26 NaHCO_3_, 1.2 NaH_2_PO_4_, 10 glucose) bubbled with 95% oxygen, 5% carbon dioxide. Cut slices were transferred to artificial cerebrospinal fluid (aCSF, in mM: 124 NaCl, 4.5 KCl, 2 CaCl_2_, 1 MgCl_2_, 26 NaHCO_3_, 1.2 NaH_2_PO_4_, 10 glucose) bubbled with 95% oxygen, 5% carbon dioxide and incubated at 32°C for 30 min for recovery and stored at room temperature until recording.

### Whole-cell patch-clamp recording

Slices were hemisected and placed into a recording chamber perfused with temperature controlled aCSF (28-31°C) bubbled with 95% oxygen, 5% carbon dioxide. VGLUT2+ and VGLUT2-cells were identified under 40x magnification by mCherry fluorescence. Recordings were performed using a borosilicate glass pipette (3-5MΩ resistance) attached to a headstage amplifier (Axon Instruments CV-7B). Current- and voltage-clamp was performed using a Multiclamp 700B amplifier (Axon CNS) and signals were filtered at 2kHz, digitized at 10kHz using the Digidata 1440A A/D converter (Axon CNS) and acquired using the Clampex 10.4.1.4 software (Molecular Devices).

Type 1 neurons were identified by a <145pF membrane capacitance, by exhibiting a depolarization block during a 500ms depolarizing current step, and by a voltage-gated calcium current <1000pA. Type 2 neurons were identified by a >145pF membrane capacitance, burst firing, and a voltage-gated calcium current >1000pA [4,37].

Neurons with membrane capacitance <100PF were identified as Type 1 and those with >145pF membrane capacitance were identified as Type 2. For neurons with membrane capacitance 100pF-145pF, Type 2 neurons were identified by burst firing behavior and the absence of depolarization block, while Type 1 neurons lacked these features. During voltage clamp recordings, when action potentials cannot be induced, Type 2 neurons were identified by a voltage-gated calcium current >1000pA.

Intrinsic excitability and ACC responsivity were recorded in current-clamp configuration at resting membrane potentials using a potassium gluconate-based internal solution (in mM: 126 potassium gluconate, 4 KCl, 10 HEPES, 4 Mg-ATP, 0.3 Na-GTP, 10 Phosphocreatine). Optically evoked excitatory postsynaptic currents (EPSC) were recorded in voltage-clamp configuration using a CsMeSO_3_-based internal solution (in mM: 120 CsMeSO_3_, 10 HEPES, 5 NaCl, 10 TEA-Cl, 0.3 Na-GTP, 4 Mg-ATP, 5 QX-314). To isolate monosynaptic EPSCs, 1μM Tetrodotoxin and 100μM 4-amino pyridine were added to the aCSF. ACC-evoked release of asynchronous EPSCs were recorded in voltage-clamp configuration in a strontium-containing aCSF (in mM: 124 NaCl, 4.5 KCl, 2 SrCl_2_, 1 MgCl_2_, 26 NaHCO_3_, 1.2 NaH_2_PO_4_, 10 glucose). ACC terminals were optically stimulated with 470nm light from an optic fiber placed approximately 150μm from the recording electrode.

Passive membrane properties (membrane capacitance, membrane resistance and access resistance) were recorded in voltage-clamp configuration using a 5mV depolarizing step. In current-clamp configuration, burst properties were determined by injecting a 2-5ms long 1-3nA current. Intrinsic excitability was measured by ramping up current at a rate of 0.5pA/ms for 800ms. The threshold potential of the first evoked action potential was measured along with the current required to generate the action potential (rheobase). Firing rate was measured by injecting a 500ms step current in 40pA steps from -100pA to 300pA. Three 5ms light-pulses were delivered with a 150ms inter-stimulus interval at intensities ranging from 0mW to 3mW in 0.6mW steps. The area under the response was measured using the 200ms prior to the first stimulus as baseline. The number of generated action potentials were counted and divided by the number of stimulation pulses to measure the firing fidelity.

Recordings from step-wise current injections were used to analyze action potential dynamics using custom Matlab scripts. The threshold potential, peak, width (half-way between the threshold and peak), and after-hyperpolarization (the difference in voltage between the threshold potential and the lowest membrane potential immediately following the action potential) was calculated for each action potential. The values from the first action potential in each neuron were extracted to determine the dynamics of the first evoked action potential. The values from the current step resulting in the maximum firing rate were averaged for each neuron. The maximum depolarization rate and repolarization rate was calculated by finding the maximum and minimum value of the differential of the action potential. For the maximum firing rate action potential the membrane potential from -2 to 4 ms from the peak of each action potential was first averaged to create an average action potential at the maximum firing rate. Depolarization and repolarization rate was then calculated based on the differential of the average action potential.

For voltage-clamp recordings, voltage-gated calcium currents were recorded while holding the cell at -60mV as the peak negative current generated by depolarizing the cell to -10mV for 500ms. Using a Cs-Mes-based internal solution, cells were held at -60mV and passive membrane properties were recorded with a 5mV depolarizing step. Paired-pulse recordings were obtained while holding the cell at -60mV and stimulating with two 1ms light pulses with a 50ms inter-stimulus interval at intensities ranging from 0.3mW to 1.5mW. The peak amplitude of the first EPSC was used to generate an input-output (I-O) curve and the paired-pulse ratio was measured from the response to 1.5mW stimulation. AMPA and NMDA currents were recorded by giving a single 1ms stimulus while holding the cell at -60mV and +40mV, respectively (5 sweeps per holding potential). The stimulation intensity was adjusted to evoke approximately -500pA peak EPSC at -60mV. AMPA current was measured at 0-20ms following stimulation and NMDA current was measured at 50ms following stimulation.

Asynchronous EPSCs were evoked with a single 1ms light pulse was given every 20s for a total of 15 sweeps. Stimulation intensity was adjusted to produce a peak EPSC of approximately 300-400pA. Asynchronous EPSC amplitude and frequency was measured for 100ms after the EPSC peak.

### Confocal imaging of viral expression

Following whole-cell patch-clamp recordings, 250uM thick coronal brain slices containing the ACC and claustrum were fixed in 4% paraformaldehyde in phosphate-buffered saline (PBS) at 4°C overnight, then were rinsed and stored in PBS until imaging. Confocal imaging was performed using a Nikon A1R + X1 spinning disk microscope. All slices were imaged at 20X magnification (NA=0.75) and analyzed using Nikon NIS-Elements software and ImageJ/Fiji.

### Data Analysis

Current clamp data was analyzed using custom scripts in MATLAB R2021A (MathWorks). Voltage-gate calcium currents and EPSCs were analyzed in Clampfit 11.2.1.00 (Molecular Devices). Asynchronous EPSC amplitude and frequency was analyzed in Minhee Analysis 1.1.3 [41]. Statistical analysis and graphing was performed in GraphPad Prism 10. Cells that were identified as fast-spiking interneurons or which did not exhibit properties consistent with Type 1 or Type 2 physiology were excluded from analysis. Iterative Grubbs outlier analysis (α=0.5) was used to identify outlier cells which were excluded from analysis. Data from included cells were subsequently averaged by animal. A second iterative Grubbs outlier analysis was performed to identify outlier animals who were excluded from analysis.

A two-sample T-test was used to compare air and ethanol exposed animals for Type 1 and Type 2 neurons. If the assumption that data conformed to a normal distribution was rejected by the Anderson-Darling test, a Mann-Whitney U test was used for comparisons. If the assumption of equal variance between groups was rejected using an F test, a Welch’s correction was performed. For comparing input-out data a 2-way repeated-measures ANOVA was used with a post-hoc Sidák correction for multiple comparisons. Sex-alcohol interactions were tested using 2-or 3-way ANOVAs with post-hoc Sidák correction for multiple comparisons. A Fisher’s exact test was used to compare proportions of Type 1 and Type 2 neurons in Air- and Ethanol-exposed mice and to determine proportions of VGLUT2 expression by physiological subtype in an equal number of randomly sampled Type 1 and Type 2 neurons.

## Results

### Chronic intermittent ethanol enhanced ACC excitation of the claustrum

The claustrum possesses two physiologically-defined (Types 1 and 2) as well as two genetically-defined - VGLUT2-positive (VGLUT2+) and VGLUT2-negative (VGLUT2-) - projection neuron subtypes that receive primary input from the anterior cingulate cortex (ACC) [3,4]. Whether these populations are overlapping is unexplored, as is their response to chronic intermittent ethanol (CIE). Thus, we exposed VGLUT2-Cre mice to chronic intermittent ethanol (CIE) vapor or air for five weeks, four nights per week, and recorded from claustrum neurons to measure the response to optically evoked inputs from the ACC three to seven days after the last vapor exposure (**Figure 1A**). Intracranial viral injections were used to express channelrhodopsin2 (ChR2) in the ACC and mCherry cre-dependently in claustrum. This allowed claustrum identification (**Figure 1B**) [35,39,42,43]. We recorded from Type 1 and Type 2 projection neurons, which exhibit burst firing and membrane capacitance differences (**Figure 1C**) [37]. CIE did not affect the proportion of Type 1 and Type 2 neurons (**Figure 1D**; Fisher’s exact test, p = 0.3). Claustrum neurons from CIE-exposed mice had significantly lower membrane capacitance independent of subtype (**Figure 1E**, 2-way ANOVA, F[1,68]=9.86, p=0.003) and Type 2 neurons exhibited larger membrane capacitance than Type 1 neurons (2-way ANOVA, F[1,68]=173.8, p<0.0001) following air (Šídák’s multiple comparisons test, t[68]=9.446, p<0.0001) and CIE treatment (Šídák’s multiple comparisons test, t[68]=9.203, p<0.0001).

**Figure1:**
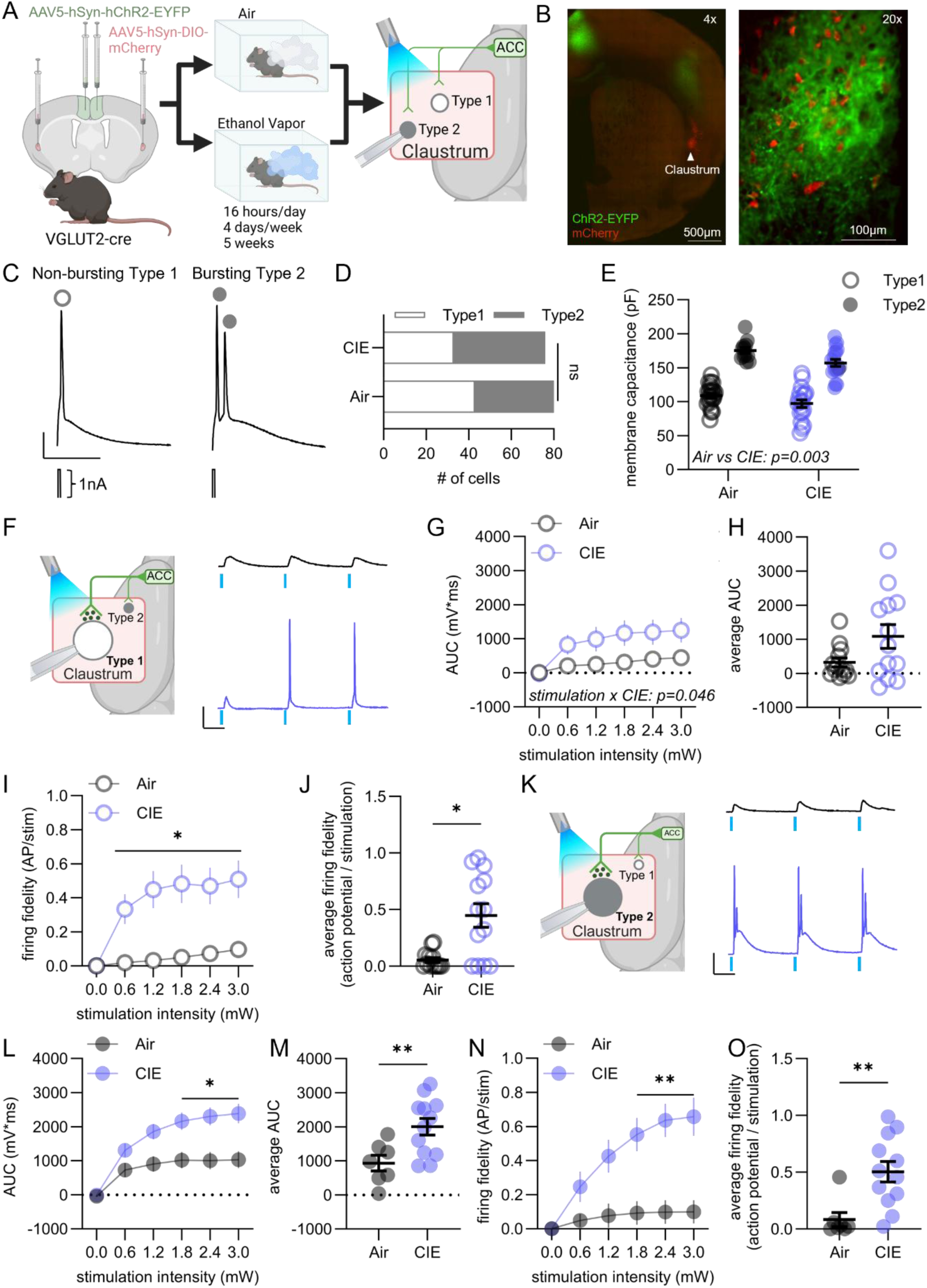
Chronic intermittent ethanol (CIE) enhanced anterior cingulate cortex (ACC) synaptic drive of Type 1 and Type 2 claustrum projection neurons. A) Schematic of the experimental design. B) Channelrhodopsin 2 (ChR2)-EYFP expression in the ACC and terminal labeling in the claustrum (right), along with VGLUT2-cre driven expression of mCherry in claustrum neurons. Insert; Overlapping mCherry and ChR2-EYFP-expression in claustrum. C) Example traces of Type 1 neurons (non-bursting) and Type 2 neurons (bursting) in response to a brief current injection (1 nA, 2 ms). D) From a random sample, a similar number were defined as Type 1 and Type 2 neurons following air and CIE. E) Type 2 neurons had significantly greater membrane capacitance than Type 1 neurons after both air and CIE. CIE reduced membrane capacitance independently from subtype. F) When activating ACC afferents and recording from Type 1 neurons (Air: 44 cells from N= 13 animals, Ethanol: 33 cells from N = 13 animals), CIE increased postsynaptic responses, including G) the area under the curve (AUC) of the voltage response with increasing light stimulation intensities, but H) did not alter the average AUC across stimulation intensities. I) CIE increased the action potential (AP) firing fidelity in response to ACC input activation as well as J) the average firing fidelity across stimulation intensities in Type 1 neurons. K) When activating ACC afferents and recording from Type 2 neurons (Air: 14 cells from N = 7 animals, EtOH: 23 cells from N = 12 animals), CIE increased postsynaptic responses including L) the AUC at high stimulation intensities, M) the average AUC, N) firing fidelity at higher light stimulation intensities and O) the average firing fidelity. Mean +/- SEM. Individual datapoints = mouse average. *p<0.05, **p<0.01, ***p<0.001, ****p<0.0001. Scale bars: 20 mV (vertical), 50 ms (horizontal).

When recording responses to increasing intensities of light stimulation in Type 1 neurons (**Figure 1F**), we observed a significant interaction between light intensity and treatment (**Figure 1G**; 2-way RM ANOVA, F[1.220,29.27]=4.039, p=0.046) in the area under the curve (AUC) of the voltage response but no significant difference between air- and CIE-exposed mice was observed at any individual light intensity or when averaging across light intensities (**Figure 1H**). A significant interaction between light intensity and CIE-treatment was also observed for the firing fidelity of Type 1 claustrum neurons in response to ACC activation (**Figure 1I**; 2-way RM ANOVA, F[1.626, 39.02]=10.42, p=0.0005) with a post-hoc test revealing a significantly higher firing fidelity at each individual light intensity (Šídák’s multiple comparisons test, 0.6mW t[12.64]=3.501, p=0.02, 1.2mW t[12.68]=3.787, p=0.01, 1.8mW t[13.54]=3.716, p=0.01, 2.4mW t[14.35]=3.499, p=0.02, 3.0mW t[14.84]=3.513, p=0.02), and when averaging across light intensities (**Figure 1J**; Mann Whitney test, Mann-Whitney U=38, p=0.01). The effect of CIE was not significantly affected by sex for either AUC or firing fidelity (**Suppl. Figure S1B-C**).

Similar evidence of a hyper-excitatory drive by ACC following CIE was observed when recording from Type 2 claustrum neurons (**Figure 1K**). We found a significant interaction between light intensity and CIE treatment on AUC of the response in Type 2 neurons (**Figure 1L**; 2-way RM ANOVA, F[1.586,26.96]=8.185, p=0.003) with a post-hoc test revealing a significant increase in the AUC of CIE-exposed mice at greater light intensities (Šídák’s multiple comparisons test, 0.6mW t[15.55]=1.967, p=0.3, 1.2mW t[15.64]=2.967, p=0.05, 1.8mW t[16.11]=3.259, p=0.03, 2.4mW t[16.04]=3.536, p=0.02, 3.0mW t[16.70]=3.569, p=0.01). The AUC averaged across all light intensities was significantly greater in Type 2 neurons from CIE-exposed mice (**Figure 1M**; t-test, t[17]=2.936, p=0.009). Measuring firing fidelity in Type 2 neurons following activation of ACC terminals additionally revealed a significant interaction of light intensity and CIE-treatment (**Figure 1N**; 2-way RM ANOVA, F[2.349,39.94]=7.975, p=0.0007) with greater light intensity resulting in significantly higher firing fidelity in CIE-exposed mice (Šídák’s multiple comparisons test, 0.6mW t[14.59]=2.035, p=0.3, 1.2mW t[17.00]=2.908, p=0.06, 1.8mW t[16.98]=3.849, p=0.008, 2.4mW t[16.97]=4.495, p=0.002, 3.0mW t[16.52]=4.250, p=0.003) and when averaging across all light intensities (**Figure 1O**; Mann Whitney test, Mann-Whitney U=8, p=0.002). Sex did not influence the effect of CIE on AUC or firing fidelity (**Suppl. Figure S1E-F**).

### CIE induced intrinsic hyper-excitability in Type 1 claustrum neurons

To determine if the increased ACC drive of claustrum neurons was mediated by changes in intrinsic excitability, we recorded from Type 1 (**Figure 2A**) and Type 2 claustrum neurons (**Figure 2H**) and injected a ramped current to determine the minimum current required to induce an action potential (rheobase). Type 1 neurons from CIE-exposed mice exhibited a lower rheobase (**Figure 2B**; Mann Whitney test, Mann-Whitney U=115, p=0.008), which was associated with a significantly lower threshold potential (**Figure 2C**; t-test, t[40]=3.001, p=0.005). While sex did not influence the effect of CIE on the rheobase (**Suppl. Figure S2B**), a trending interaction was observed for the threshold potential (**Suppl. Figure S2C**; 2-way ANOVA, F[1,38]=3.235, p=0.08). Type 1 claustrum projection neurons from CIE-exposed mice further exhibited a trending increase in input resistance (**Figure 2D**; t-test, t[40]=1.882, p=0.07), which was unaffected by sex (**Suppl. Figure S2D**). When injecting a step-wise current into Type 1 neurons (**Figure 2E**), a significantly greater number of action potentials was observed following CIE treatment (**Figure 2F**; 2-way RM ANOVA, F[1,40]=6.513, p=0.01) resulting in a greater maximum firing rate (**Figure 2G**; t-test, t[40]=2.186, p=0.03). Interestingly, we found a significant interaction of sex and CIE treatment on the number of action potentials (**Suppl. Figure S2E**; 3-way RM ANOVA, F[2.327,88.43]=4.512, p=0.01) and the maximum firing rate (**Suppl. Figure S2F**; 2-way ANOVA, F[1,38]=7.239, p=0.01) with CIE significantly increasing the maximum firing rate in male mice only (Šídák’s multiple comparisons test, t[38]=3.617, p=0.003); they also had lower maximum firing than females in the air-exposed controls (Šídák’s multiple comparisons test, t[38]=3.131, p=0.01). Thus, CIE increased the intrinsic excitability of Type 1 neurons.

**Figure 2:**
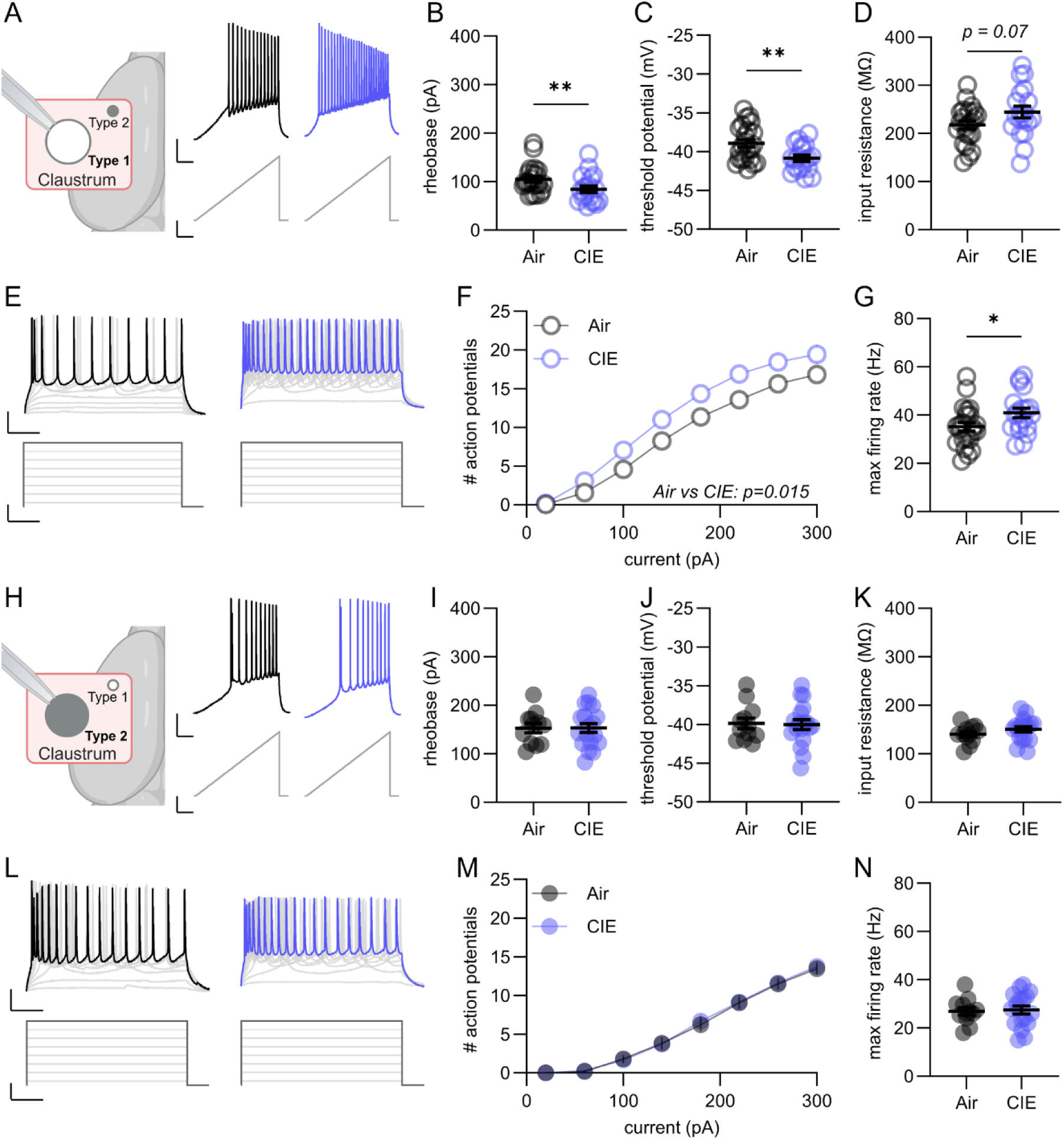
CIE increased intrinsic excitability in Type 1, but not Type 2, claustrum projection neurons. A) When recording from Type 1 neurons (Air: 54 cells from N = 23 animals, Ethanol: 45 cells from N = 19 animals) and injecting a ramped current, CIE significantly increased the excitability, including B) reducing the rheobase, C) lowering the threshold potential but D) only resulted in a trending increase in the input resistance. E) Upon step-wise current injections into Type 1 neurons, CIE F) increased the number of action potentials at increasing current amplitudes resulting in G) greater maximum firing rate compared to Type 1 neurons from air-exposed mice. H) When recording from Type 2 neurons (Air: 28 cells from 12 animals, Ethanol: 43 cells from 18 animals) while injecting a ramped current, no effect of CIE was observed, including in the I) rheobase, J) threshold potential, or K) input resistance. L) Similarly, when injecting a step-wise current no effect of CIE was observed on the M) number of action potentials over increasing current amplitudes or N) the maximum firing rate. Mean +/- SEM. Individual datapoints = mouse averages. *p<0.05, **p<0.01, ***p<0.001, ****p<0.0001. Scale bars: 20mV or 50 pA (vertical), 100 ms (horizontal).

In contrast to Type 1 neurons, Type 2 claustrum projection neurons from CIE-exposed mice exhibited similar rheobase, threshold potential, and input resistance to air-exposed mice (**Figure 2I-K**). While we did not observe any effect of sex on rheobase (**Suppl. Figure S2H)** or input resistance (**Suppl. Figure S2J**), Type 2 neurons from male mice exhibited a significantly elevated threshold potential independent of CIE treatment (**Suppl. Figure S2I**; 2-way ANOVA, F[1,26]=7.508, p=0.01). Further, when injecting a step-wise current into Type 2 neurons (**Figure 2L**), no difference in the observed number of action potentials or maximum firing rate was found between air- and CIE-exposed mice (**Figure 2M-N**). However, Type 2 neurons from male mice, compared to females, fired significantly fewer action potentials (**Suppl. Figure S2K**; 3-way RM ANOVA, F[1.549,40.28]=3.911, p=0.04) and a trending lower maximum firing rate (**Suppl. Figure S2L**; 2-way ANOVA, F[1,26]=3.739, p=0.06) independent of CIE treatment.

### CIE altered action potential dynamics in both Type 1 and Type 2 neurons

To better understand the underlying physiological mechanisms allowing for the enhanced firing of Type 1 claustrum neurons, we analyzed the dynamics of the averaged action potential elicited by the current step resulting in the maximum firing rate (**Figure 3A**). The phase plot derived from the action potential reveals small differences between air- and CIE-exposed mice (**Figure 3B**). Surprisingly, CIE did not seem to affect the threshold potential during the maximum firing rate in Type 1 neurons (**Figure 3C**) but did result in a trending reduction in the peak of the action potential (**Figure 3D**; t-test, t[40]=1.919, p=0.06). In a similar analysis of the first action potential produced during the injection of the minimum current step required to elicit firing, we did observe that CIE treatment lowered the threshold potential (**Suppl. Figure S4C**; t-test, t[40]=2.538, p=0.02) but no difference was observed in the amplitude of action potential peak (**Suppl. Figure S4D**). While we did not observe any difference in the width of the action potential (**Figure 3E**), CIE significantly reduced the size of the after-hyperpolarization (**Figure 3F**; Welch’s test, t[33.39]=2.670, p=0.01). This suggests that the increased firing rate of Type 1 neurons following CIE treatment was due to a shorter refractory period. This was further supported by our finding that no effect of CIE was found on the action potential maximum depolarization or repolarization rate (**Figure 3G-H**). We did not find that sex had any impact on Type 1 action potential dynamics (**Suppl. Figure S3A-G**). For the first encountered action potential, we did not observe an effect of CIE on either the action potential width, after-hyperpolarization, depolarization, or repolarization (**Suppl. Figure S4E-H**).

**Figure 3:**
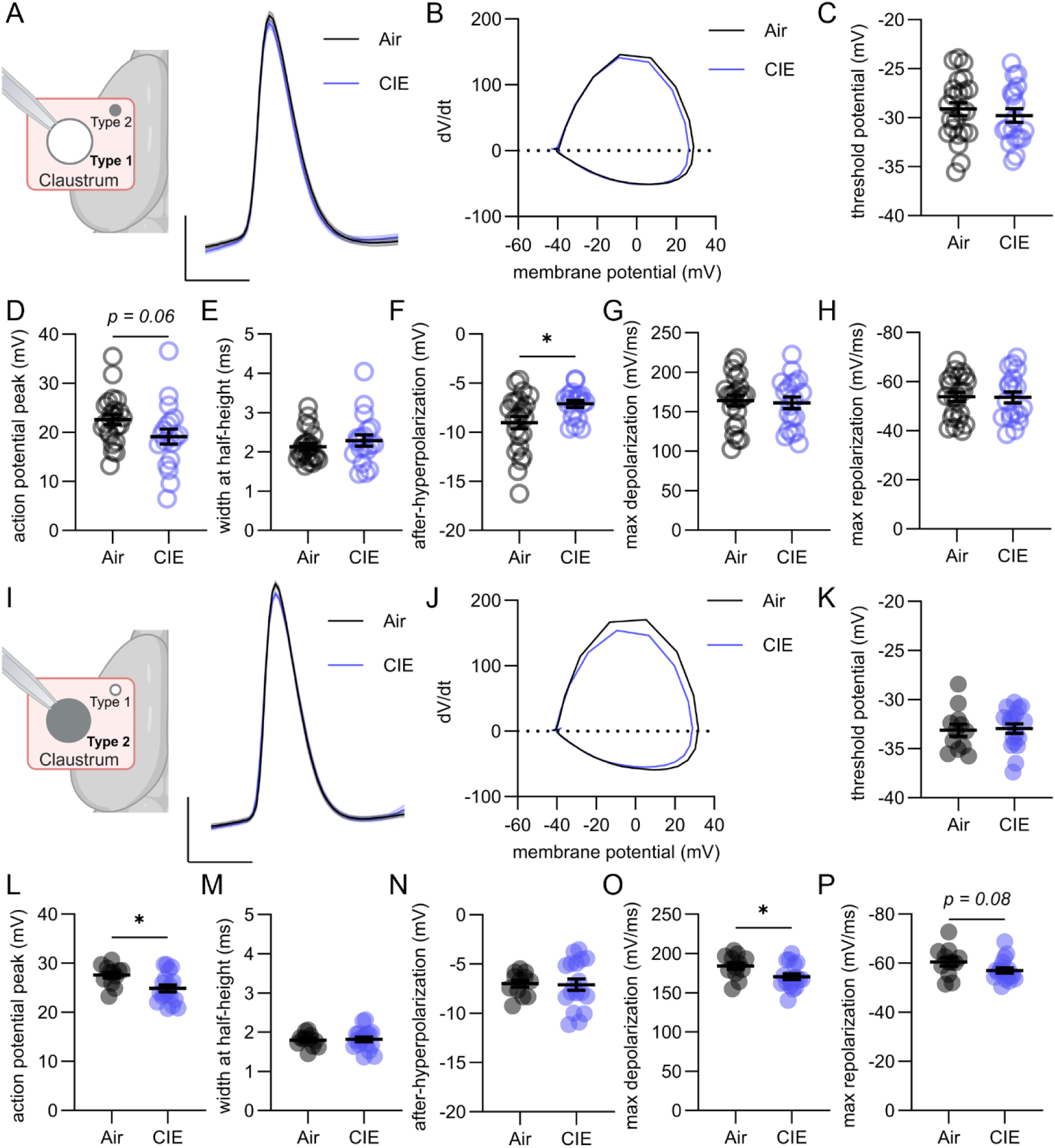
CIE caused distinct action potential changes in Type 1 and Type 2 claustrum projection neurons. A) The action potentials recorded during the current step resulting in maximum firing rate were averaged and dynamics compared between Type 1 neurons (Air: 54 cells from N = 23 animals, Ethanol: 45 cells from N = 19 animals) in air- and CIE-exposed mice. B) Phase plot of averaged action potential from air and CIE-exposed mice. CIE did not affect the C) threshold potential but D) did result in a trending decrease in the peak amplitude of the action potential. E) CIE further did not affect the width of the action potential but F) significantly reduce the after-hyperpolarization. CIE did not affect either the G) depolarization or the H) repolarization of the action potential in Type 1 claustrum neurons I) The dynamics of the average action potential recorded during the maximum firing rate were compared between Type 2 neurons (Air: 28 cells from N = 12 animals, Ethanol: 43 cells from N = 18 animals) from air- and CIE-exposed mice. J) Phase plot of averaged action potentials in Type 2 neurons from air- and CIE-exposed mice. While CIE K) did not affect the threshold potential, L) it significantly reduced the peak of the average action potential. CIE did not affect either M) the width of the action potential or N) the after-hyperpolarization. However, CIE O) significantly reduced the depolarization rate and P) produced a trending reduction in the repolarization rate of the average action potential. Mean +/- SEM. Individual data points = mouse average. *p<0.05, **p<0.01, ***p<0.001, ****p<0.0001. Scale bars: 20 mV (vertical), 2 ms (horizontal).

To determine if CIE induced changes to the action potential dynamics of Type 2 neurons, we performed the same analysis (**Figure 3I**). Surprisingly, the phase plot suggested greater differences in the action potential dynamics of Type 2 neurons following CIE treatment compared to what was observed in Type 1 neurons (**Figure 3J**). Like Type 1 neurons, the threshold potential of Type 2 neurons was not affected by CIE (**Figure 3K**) but the action potential peak was significantly reduced in CIE-exposed mice (**Figure 3L**; t-test, t[28]=2.697, p=0.01). Unlike Type 1 neurons, Type 2 neurons did not exhibit any difference in either action potential width or after-hyperpolarization between air- and CIE-exposed mice (**Figure 3M-N**). When analyzing the first action potential, CIE significantly reduced the peak but did not affect threshold potential, width, or after-hyperpolarization (**Suppl. Figure S4K-N**; t-test, t[28]=2.458, p=0.02). Surprisingly, CIE reduced the action potential maximum depolarization rate (**Figure 3O**; t-test, t[28]=2.415, p=0.02) and resulted in a trending decrease in the maximum repolarization rate (**Figure 3P**; t-test, t[28]=1.844, p=0.08) of Type 2 neurons. Analysis of the first action potential similarly showed a trending increase in the depolarization in CIE-exposed mice while no effect was observed on the repolarization (**Suppl. Figure S4O-P**; Mann Whitney test, Mann-Whitney U=64, p=0.0649).

Further, the Type 2 neurons also exhibited an elevated threshold potential in male mice (**Suppl. Figure S3I**; 2-way ANOVA, F[1,26]=4.931, p=0.04), while sex did not affect other dynamics (**Suppl. Figure S3J-N**).

### CIE strengthened ACC synaptic transmission onto Type 2 neurons

Synaptic mechanisms of CIE-induced enhancement of claustrum response to ACC inputs were tested by recording evoked excitatory postsynaptic currents (EPSCs) to optical stimulation of ACC terminals in Type 1 (**Figure 4A**) and Type 2 neurons (**Figure 4I**). Type 1 neurons from air-and CIE-exposed mice exhibited similar EPSC amplitudes (**Figure 4B**) and CIE did not affect the paired-pulse ratio (50 ms inter-pulse interval) of optically evoked EPSCs (**Figure 4C**) but surprisingly increased the AMPA:NMDA ratio (**Figure 4D-E**; Welch’s test, t[6.790]=2.475, p=0.04). However, recordings of asynchronously released EPSCs (aEPSCs) using strontium-containing aCSF **(Figure 4F)** revealed no difference in either the frequency (**Figure 4G**) or amplitude (**Figure 4H**) of aEPSCs between Type 1 neurons from air- or CIE-exposed mice. While we did observe a trending interaction between sex, CIE treatment, and the light stimulation intensity on EPSC amplitude (**Suppl. Figure S5B**; 3-way RM ANOVA, F[1.198,13.18]=3.115, p=0.1), we found no other evidence of sex influencing synaptic transmission onto Type 1 neurons (**Suppl. Figure S5C-F**).

**Figure 4:**
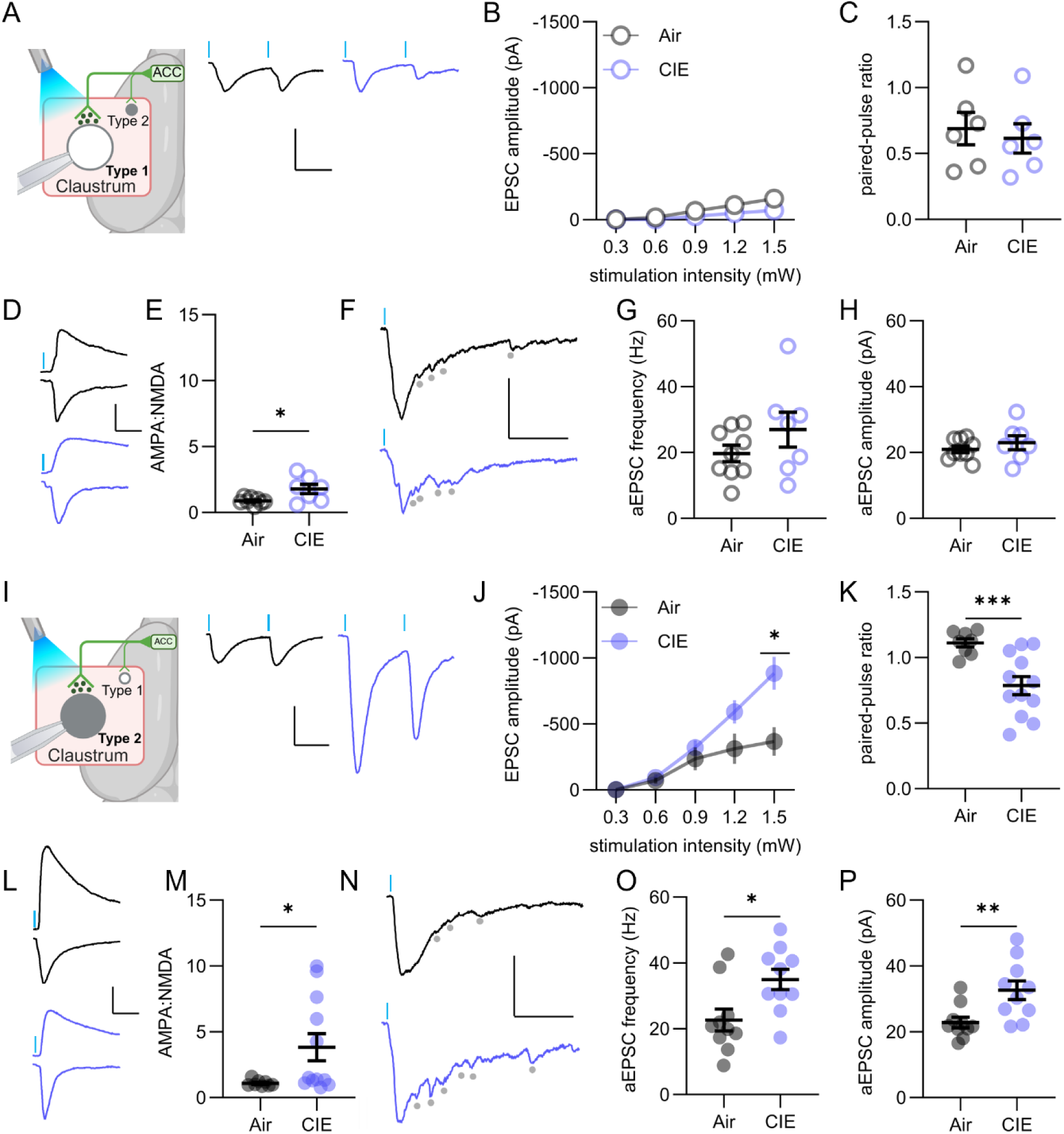
CIE strengthened ACC synapses onto Type 2, but not Type 1, claustrum projection neurons through pre- and post-synaptic mechanisms. A) When activating ACC afferents and recording EPSCs in Type 1 neurons (Air: 19 cells from N = 9 animals, Ethanol: 17 cells from N = 7 animals) no effect of CIE was observed on either B) the amplitude at increasing stimulation intensities or C) the paired-pulse ratio. D) However, when recording the AMPA- and NMDA-mediated current at -60 mV and +40 mV, respectively E) CIE significantly increased the AMPA:NMDA ratio. F) When recording asynchronously released EPSCs (aEPSC) from activated ACC afferents in Type 1 neurons (Air: 13 cells from N = 9 animals, EtOH: 10 cells from N = 7 animals), CIE did not affect either G) the frequency or H) the amplitude. I) When activating ACC-afferents and recording Type 2 claustrum neurons (Air: 23 cells from N = 9 animals, Ethanol: 28 cells from N = 12 animals) EPSCs, CIE J) significantly increased EPSC responses at larger stimulation intensities and K) significantly reduced the paired-pulse ratio. When recording the AMPA- and NMDA-mediated current at -60 mV and +40 mV, respectively, M) CIE significantly increased the AMPA:NMDA ratio. N) When activating ACC afferents and recording aEPSCs in Type 2 neurons (Air: 26 cells from N = 10 animals, Ethanol: 23 cells from N = 10 animals), CIE significantly increased both O) the frequency and P) the amplitude. Mean +/- SEM. Individual data points = mouse average. *p<0.05, **p<0.01, ***p<0.001, ****p<0.0001. Scale bars: 300 pA (vertical), 30 ms (horizontal).

In Type 2 neurons, there was a significant interaction between light intensity and ethanol exposure (**Figure 4J**; 2-way RM ANOVA, F[1.237,22.27]=7.036, p=0.01) with neurons from CIE-exposed mice exhibiting significantly larger EPSCs at 1.5mW stimulation (Šídák’s multiple comparisons test, t[17.89]=3.161, p=0.03). This was associated with a significantly lower paired-pulse ratio (**Figure 4K**; Welch’s t-test, t[15.10]=4.341, p=0.0006) and increased AMPA:NMDA ratio (**Figure 4L-M**; Mann Whitney test, Mann-Whitney U=20, p=0.03), suggesting CIE-induced pre- and post-synaptically mediated potentiation of ACC-Type 2 synapses. Neither of these effects were influenced by sex (**Suppl. Figure S5 G-J**). Recording aEPSCs (**Figure 4N**), we found CIE increased both the frequency (**Figure 4O**; t-test, t[18]=2.716, p=0.01) and amplitude (**Figure 4P**; t-test, t[18]=3.019, p=0.007) of aEPSCs. We observed a trending increase in aEPSC frequency (**Suppl. Figure S5K**; 2-way ANOVA, F[1,16]=3.101, p=0.1) and a significant increase in amplitude (**Suppl. Figure S5L**; 2-way ANOVA, F[1,16]=9.202, p=0.0079) in male mice with a trending interaction between sex and CIE treatment on the aEPSC amplitude (2-way ANOVA, F[1,16]=3.416, p=0.08).

### VGLUT2 molecular subtyping aligned with physiological subtyping and their alcohol effects

To determine if alcohol-effects were dependent on molecular expression, we re-typed the neurons based on VGLUT2 expression as identified by mCherry fluorescence. We found that Type 2 neurons were significantly more likely to express VGLUT2 than Type 1 neurons (**Figure 5A**; Fisher’s exact test, p=0.003) with 63% of Type 2 neurons and only 38% of Type 1 neurons expressing VGLUT2. In accordance with this finding, we observed that VGLUT2-expressing (VGLUT2+) neurons expressed higher membrane capacitance than VGLUT2-non-expressing (VGLUT2-) neurons, independently of CIE treatment (**Figure 5B**; 2-way ANOVA, F[1,79]=23.98, p<0.0001) with a trending difference in air-exposed mice (Šídák’s multiple comparisons test, t[79]=2.180, p=0.06) and a significant difference in CIE-exposed mice (Šídák’s multiple comparisons test, t[79]=4.687, p<0.0001). CIE treatment significantly reduced the membrane capacitance (2-way ANOVA, F[1,79]=6.059, p=0.02) with a trending interaction between CIE treatment and VGLUT2 expression being observed (2-way ANOVA, F[1,79]=3.560, p=0.0629).

**Figure 5:**
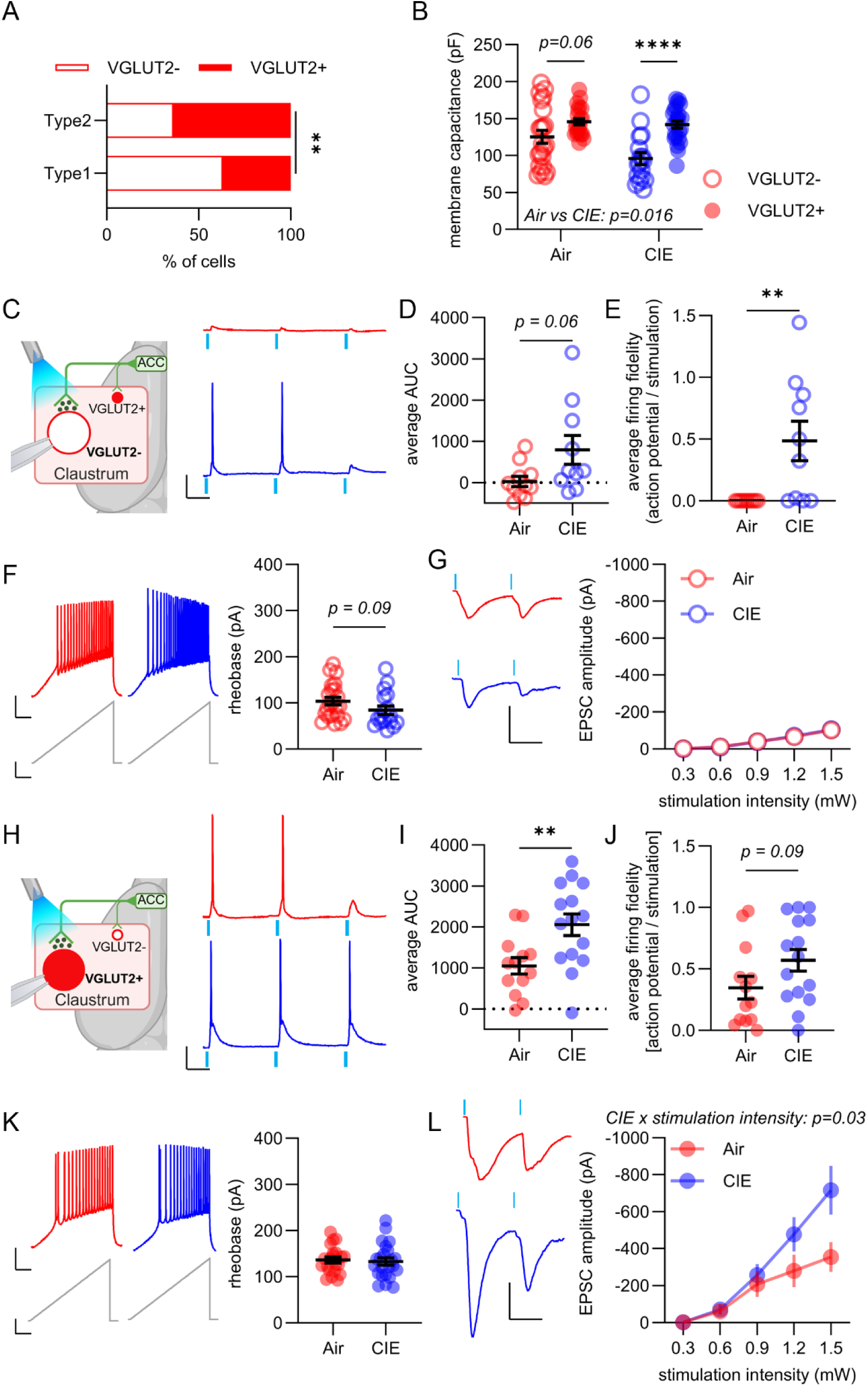
The majority of VGLUT2 neurons are Type 2 neurons and CIE affects VGLUT-negative and VGLUT2-positive neurons similarly to Type1 and Type 2 neurons, respectively. A) Type 2 neurons were more likely to express VGLUT2 than Type 1 neurons. B) VGLUT2-positive (VGLUT2+) neurons had greater membrane capacitance than VGLUT2-negative (VGLUT2-) neurons. C) When activating ACC afferents and recording the response in VGLUT2- claustrum neurons (Air: 22 cells from N = 11 animals, Ethanol: 24 cells from N = 10 animals), CIE increased the response, including D) a trending increase in the average area under the curve of the voltage response and E) a significant increase in the average action potential (AP) firing fidelity. F) When injecting a ramped current into VGLUT2- neurons (Air: 39 cells from N = 22 animals, Ethanol: 36 cells from N = 17 animals), CIE produced a trending lowering of the rheobase. G) When activating ACC afferents and recording EPSCs in VGLUT2- neurons (Air: 18 cells from N = 10 animals, Ethanol: 18 cells from N = 9 animals), CIE did not affect the amplitude over increasing light stimulation intensities. H) When activating ACC afferents and recording responses in VGLUT2+ neurons (Air: 33 cells from N = 13 animals, Ethanol: 36 cells from N = 15 animals), CIE I) significantly increased the average area under the curve (AUC) of the light-evoked response and J) produced a trending increase in the AP firing fidelity. K) When injecting a ramped current into VGLUT2+ neurons (Air: 53 cells from N = 21 animals, Ethanol: 52 cells from N = 23 animals), CIE did not affect the rheobase. L) When activating ACC afferents and recording EPSCs in VGLUT2+ neurons (Air: 33 cells from N = 12 animals, Ethanol: 30 cells from N = 12 animals), CIE significantly enhanced the EPSCs at higher stimulation intensities. Mean +/-SEM. Individual data points = mouse average. *p<0.05, **p<0.01, ***p<0.001, ****p<0.0001. Vertical scale bars: 20mV or 50pA (F and K) or 300pA (G and L). Horizontal scale bars: 50ms or 30ms (G and L).

When recording from VGLUT2- neurons, we found that ACC drive was enhanced in CIE-exposed mice (**Figure 5C**). A trending enhancement of the AUC of the response to ACC was observed in VGLUT2- neurons from CIE-exposed mice (**Figure 5D**; Welch’s test, t[11.24]=2.084, p=0.06) and the firing fidelity was significantly greater compared to air-treated mice (**Figure 5E**; Mann Whitney test, Mann-Whitney U=16.5, p=0.001). Following CIE treatment, VGLUT2-claustrum neurons exhibited a trending lower rheobase (**Figure 5F**; Mann Whitney test, Mann-Whitney U=126, p=0.0867). Additionally, when injecting a step-wise current, CIE produced a trending increase in the number of action potentials (**Suppl. Figure S6F**; 2-way RM ANOVA, F[1,37]=3.358, p=0.08) and in the maximum firing rate (**Suppl. Figure S6G**; t-test, t[37]=1.982, p=0.06). Surprisingly, however, in VGLUT2- neurons, CIE did not affect threshold potential or input resistance (**Suppl. Figure S6H-I**). Finally, in accordance with our observations in Type 1 neurons, we did not observe any difference in EPSC amplitude of VGLUT2- neurons between air- and CIE-exposed mice (**Figure 5G**), suggesting that CIE did not affect the strength of synaptic transmission from ACC onto VGLUT2- neurons. Similarly, there was no effect of CIE on the paired-pulse ratio (**Suppl. Figure S6B**) but a significant increase in the AMPA:NMDA ratio was observed in VGLUT2- neurons from CIE-exposed mice (**Suppl. Figure S6C**; t-test, t[17]=2.473, p=0.02) as was observed in Type 1 neurons. Unlike Type 1 neurons, however, VGLUT2- neurons showed a significant increase in aEPSC frequency (**Suppl. Figure S6D**; t-test, t[13]=4.559, p=0.0005) and a trending increase in aEPSC amplitude (**Suppl. Figure S6E**; Welch’s test, t[6.099]=2.348, p=0.06) following CIE treatment.

In VGLUT2+ claustrum neurons, we also found evidence of hyper-excitatory drive by ACC following CIE treatment (**Figure 5H**). VGLUT2+ neurons from CIE-exposed mice exhibited both increased AUC (**Figure 5I**; t-test, t[26]=2.959, p=0.007) and a trending increase in firing fidelity in response to ACC input activation (**Figure 5J**; t-test, t[26]=1.747, p=0.09). Like Type 2 neurons, we did not observe any effect of CIE on VGLUT2+ intrinsic excitability. No difference was observed in VGLUT2+ rheobase between air- and CIE-exposed mice (**Figure 5K**), in action potentials evoked by step-wise current injections, or the maximum firing rate (**Suppl. Figure S6O-P**). We observed that CIE treatment lowered the threshold potential of VGLUT2+ neurons while having no effect on input resistance (**Suppl. Figure S6Q-R**; t-test, t[42]=2.263, p=0.03). As in Type 2 neurons, VGLUT2+ claustrum neurons from CIE-exposed mice showed a greater increase in EPSC amplitude with higher light-stimulation intensities (**Figure 5L**; 2-way RM ANOVA, F[1.235,25.93]=4.763, p=0.03), suggesting that CIE increased the strength of synaptic transmission from ACC onto VGLUT2+ claustrum neurons. However, no difference was observed in VGLUT2+ neurons from CIE-exposed mice in either the paired-pulse ratio or the AMPA:NMDA ratio (**Suppl. Figure S6K-L**). CIE did produce a trending increase in aEPSC frequency (**Suppl. Figure S6M**; t-test, t[19]=1.824, p=0.08) and a significant increase in aEPSC amplitude (**Suppl. Figure S6N**; t-test, t[19]=2.206, p=0.04).

## Discussion

Here, we found significant enhancement of ACC drive of claustrum projection neurons. This was mediated in Type 1 and VGLUT2- neurons primarily by increases in membrane excitability while synaptic strengthening mediated the increased activation of Type 2 and VGLUT2+ neurons by ACC activation. These data establish the claustrum as a key target of the ACC that is sensitive to the effects of chronic alcohol exposure in a neuron subtype-specific manner.

Similar to the subtype-selectivity observed here, alcohol-induced hyperexcitability occurs in D1-expressing medium spiny neurons [44,45], which is associated with voltage-gated potassium channel-mediated reduction in the after-hyperpolarization [45,46]. This suggests a possible molecular mechanism in Type 1 claustrum neurons. As observed here in Type 2 neurons, alcohol-induced strengthening of excitatory transmission is observed in the striatum and the basolateral amygdala where it is expressed both pre- and post-synaptically [33,44,47], and in a cell subtype specific manner [33,44,45,48]. These data also align with the up-regulation in AMPA receptor availability following early alcohol withdrawal [49], suggesting one possible mechanism for synaptic strengthening in Type 2 claustrum neurons. CIE-enhanced ACC drive of claustrum projection neurons may thus be mediated by subtype-specific molecular mechanisms in a manner similar to other regions that significantly contribute to behavioral changes consistent with AUD.

The claustrum innervates several cortical layers but VGLUT2+ claustrum neurons, in particular, exhibit a bias towards innervating layer 5/6 parvalbumin-positive interneurons [36]. Activation of cortical parvalbumin interneurons is causally associated with the entrainment of cortical gamma oscillations and improvements in cognitive performance [50,51]. Thus, in the very least, alcohol-induced disruption of synaptic inputs to VGLUT2+/Type 2 neurons may contribute to downstream cortical disruptions in gamma oscillation-dependent cognitive processes [52]. Given that claustrum projection neurons also innervate cortical pyramidal neurons [36,53], a reasonable hypothesis is that VGUT2-/Type1 neurons may, in part, innervate this principal neuron class. If confirmed by further investigation, it is possible that chronic alcohol may perturb claustrum control of multiple neuronal subtypes in the cortex to further disrupt cognition.

Type 1 and Type 2 claustrum neurons receive cortical excitatory afferents from frontal, parietal, sensorimotor, and limbic regions. In turn, the claustrum, including both projection neuron subtypes, projects to virtually the entire cortex [4,37,54,55], and projection-specific sub-populations of claustrum neurons are largely non-overlapping [56–59]. Despite the widespread cortical input to the claustrum and widespread output from the claustrum back to cortex, not all cortical regions project onto claustrum neurons that project back to all cortical areas. Rather, distinct patterns of cortico-claustro-cortical loops exist [60] with some loops exhibiting stronger synaptic strength than others [4]. Particularly strong are ACC excitatory inputs that synapse upon claustrum projection neurons (including Types 1 and 2) that, in turn, project back to the ACC, prelimbic prefrontal cortex, as well as parietal association and visual cortices [4]. As these regions all exhibit disrupted activity following chronic alcohol exposure [22–24,61–64], the current findings may point to claustrum as a potential node mediating common, dysregulated cortical activity in AUD that may manifest as cognitive network disruption [65,66].

Some limitations of this study are worth noting. In this study we used CIE to model cycles of binge intoxication and withdrawal in mice. While this is a useful model due to its ability to produce consistent blood ethanol concentrations in the 100-200 mg/dL range [40] and to accurately model transcriptional changes observed in human AUD [67], it does not model the actions of seeking and drinking, which invariably involve cognitive control of action (or lack thereof). Further studies are needed to determine if and how claustrum activity is affected by voluntary alcohol consumption. We also did not probe the effects of alcohol on projection-specific claustrum neuron subpopulations. This is of particular interest given that certain claustrum functions are hypothesized to rely upon these unique projection targets [5,10,39,68]. Lastly, the claustrum possesses several interneuron populations, including parvalbumin, somatostatin, and vasoactive intestinal peptide neurons that were not examined here [38].

Given the sensitivity to alcohol of several interneuron populations that govern the output of principal projection neurons across brain nuclei [40,69–71], this is an important consideration that requires future attention. Despite these important limitations, this study demonstrates that CIE disrupts ACC drive of both molecularly and physiologically defined claustrum projection neuron subtypes. This sets the stage for a deeper understanding of how cognitive control, as well as the alternative myriad claustrum functional roles, contributes to AUD pathology [5,18,72–75].

## Data Availability

All data supporting the findings of this study are available upon request from the corresponding author.

## Author Contributions

Conceptualization, A.B.W, B.N.M.; Surgical Procedures, A.B.W., S.H.S.; Histology and Microscopy, E.A.D; Electrophysiology, A.B.W.; Data Analysis, A.B.W, E.A.D.; Writing (Original Draft), A.B.W., B.N.M.; Review & Editing, A.B.W, B.N.M.; Supervision: B.N.M.; Project Administration, B.N.M.; Funding Acquisition, A.B.W. and B.N.M.

