## Supplemental Figures for "Anterior cingulate cortex activation of claustrum projection neuron subtypes is enhanced by alcohol"

**Supplemental Material**

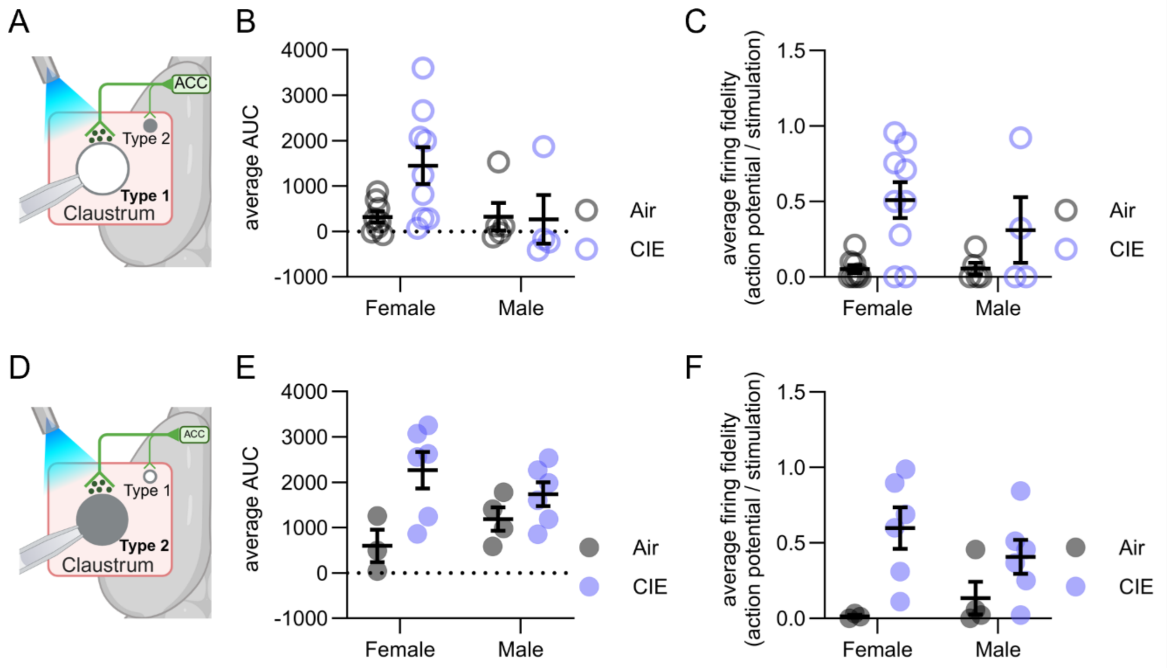


**Supplemental Figure 1: CIE sex-independently affected ACC excitatory drive of claustrum neurons.** A) When activating ACC afferents and recording responses in Type 1 claustrum neurons (Air: 18 cells from N = 8 females and 14 cells from N = 5 males, Ethanol: 20 cells from N = 9 females and 13 cells from N = 4 males), sex had no effect on either B) the average AUC of the light-evoked response or C) the AP firing fidelity in response to the ACC inputs. D) When recording responses in Type 2 neurons (Air: 7 cells from N = 3 females and 7 cells from N = 4 males, Ethanol: 13 cells from N = 6 females and 15 cells from N = 6 males), sex had no effect on either E) the AUC of the light-evoked response or F) the AP firing fidelity in response to ACC inputs. Mean +/- SEM. Individual data points = mouse average.

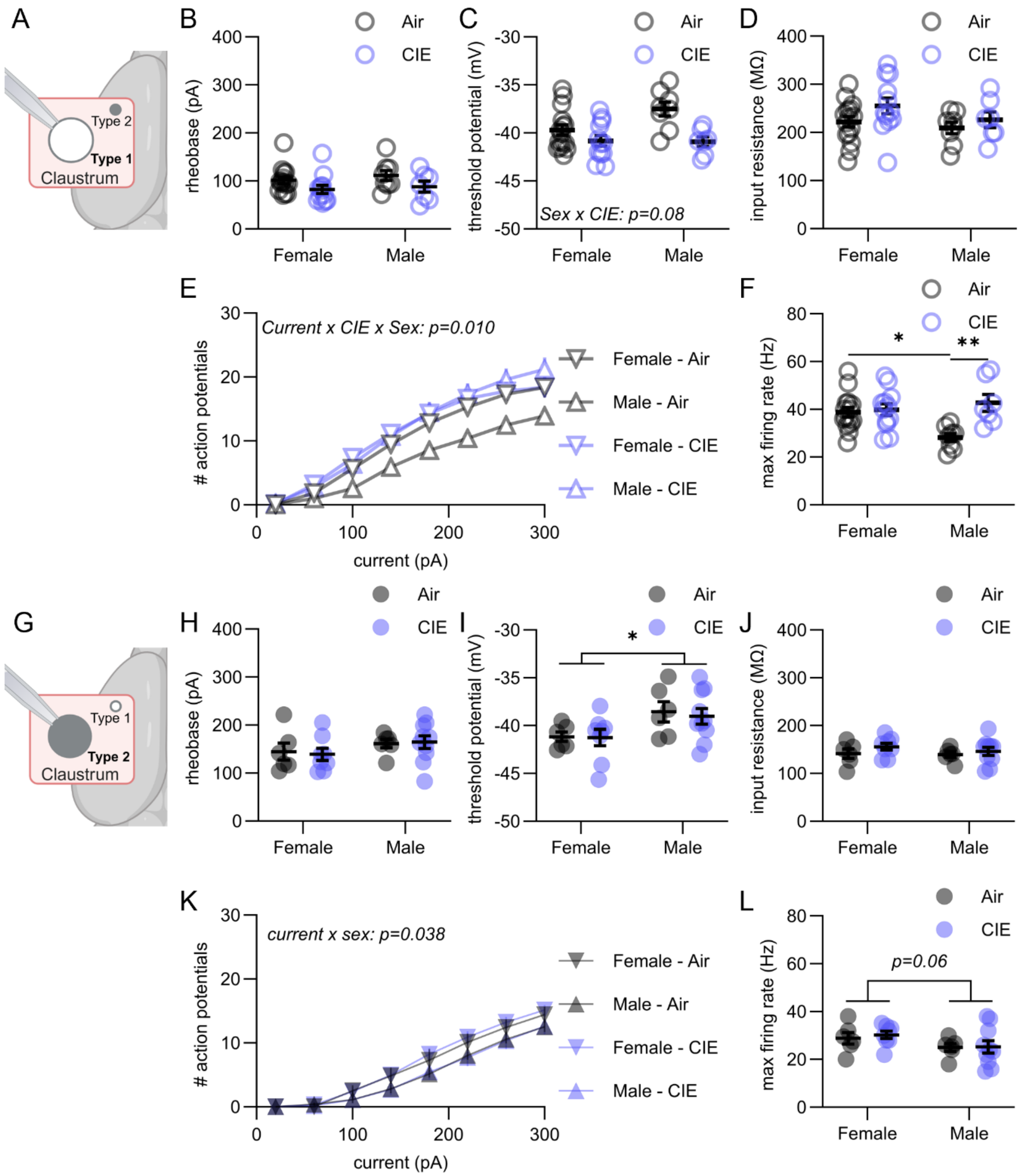


**Supplemental Figure 2: Sex-dependent CIE effects on intrinsic excitability.** A) When injecting a ramp-current into Type 1 claustrum neurons (Air: 35 cells from N = 15 females and 19 cells from N = 8 males, Ethanol: 26 cells from N = 12 females and 19 cells from N = 7 males), sex had no influence on B) the rheobase. C) While a trending influence of sex on the impact of CIE on the threshold potential was observed, D) sex did not impact the input resistance of Type 1 neurons. E) When injecting step-wise currents, sex significantly modulated the impact of CIE on the AP firing with increasing current amplitudes such that F) Type 1 neurons from air-exposed male mice had significantly lower maximum firing rate than air-exposed female mice or CIE-exposed male mice. G) When injecting a ramped current into Type 2 claustrum neurons (Air: 15 cells from N = 6 females and 13 cells from N = 6 males, Ethanol: 20 cells from N = 8 females and 23 cells from N = 10 males), H) sex had no impact on the rheobase but I) males had a significantly elevated threshold potential compared to females independent of CIE exposure. J) Sex also had no influence on the input resistance. K) When injecting a step-wise current, Type 2 neurons in male mice fired significantly fewer action potentials resulting in L) a trending lower maximum firing frequency in male mice. Mean +/- SEM. Individual data points = mouse average. *$p<0.05$, **$p<0.01$, ***$p<0.001$, ****$p<0.0001$.

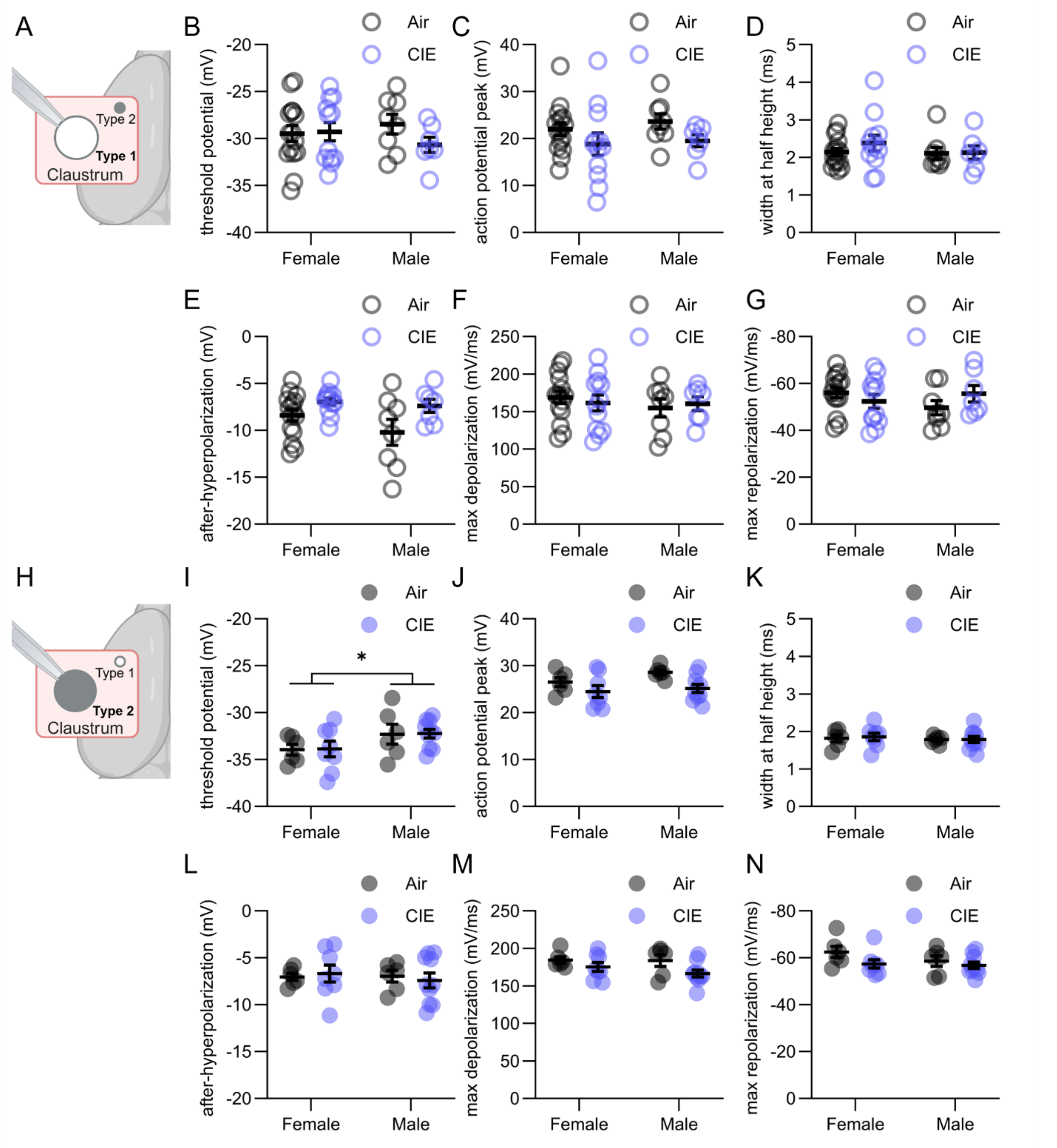


**Supplemental Figure 3: CIE sex-independently affected AP properties in Type 1 and 2 claustrum projection neurons at peak firing frequency.** A) When analyzing the dynamics of the average AP during maximum firing of Type 1 neurons (Air: 35 cells from N = 15 females and 19 cells from N = 8 males, Ethanol: 26 cells from N = 12 females and 19 cells from N = 7 males), sex had no influence on B) the threshold potential, C) the peak or D) the width of the AP, E) the after-hyperpolarization or the F) depolarization or G) repolarization of the AP. H) When analyzing the dynamics of the average AP during maximum firing of Type 2 neurons Air: 15 cells from N = 6 females and 13 cells from N = 6 males, Ethanol: 20 cells from N = 8 females and 23 cells from N = 10 males), I) male mice had significantly elevated threshold compared to females, independent of CIE exposure but sex had no impact on the AP J) peak, K) width, L) after-hyperpolarization, the M) depolarization or N) repolarization. Mean +/- SEM. Individual data points = mouse average. *$p<0.05$, **$p<0.01$, ***$p<0.001$, ****$p<0.0001$.

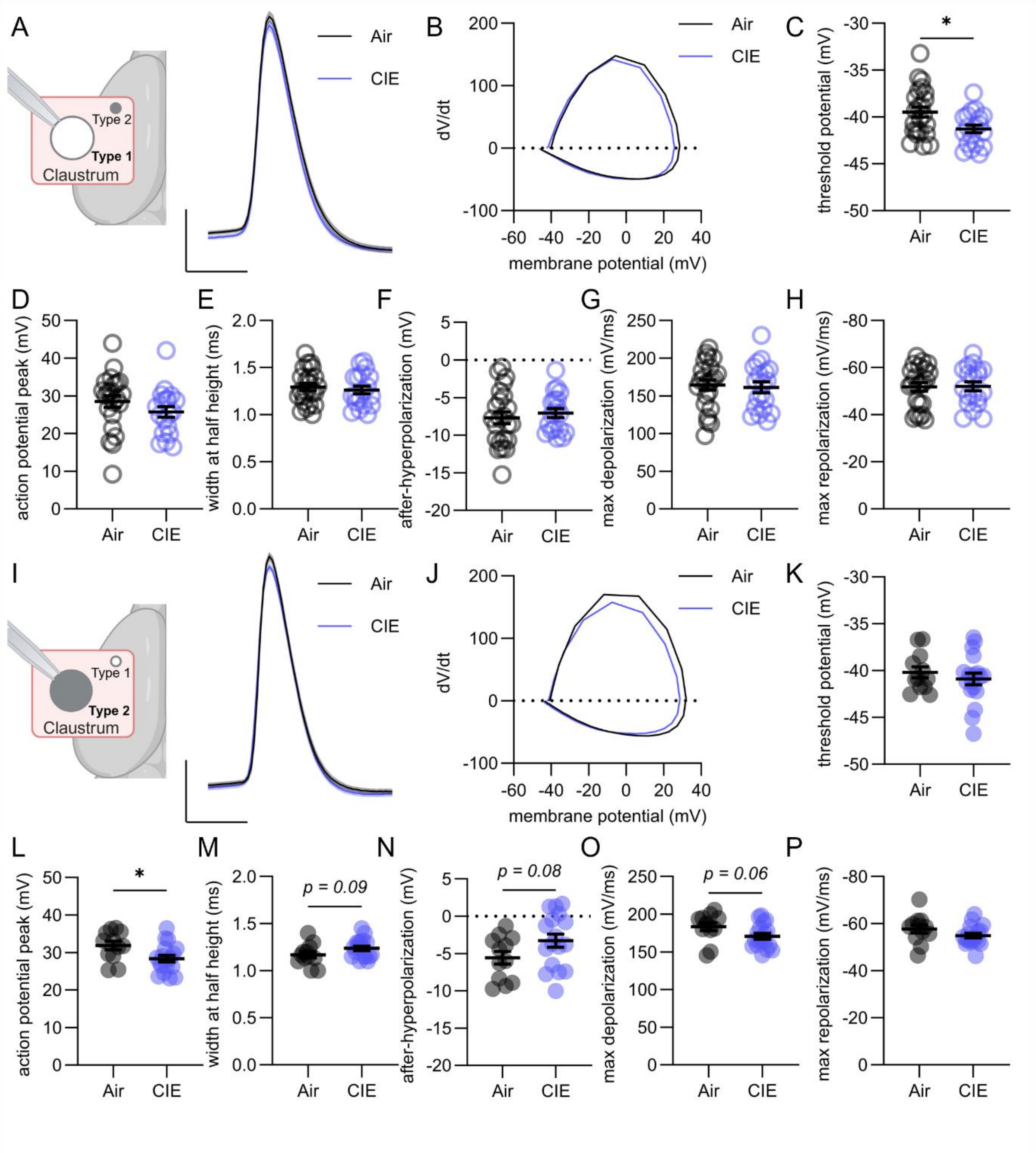


**Supplemental Figure 4: CIE differentially affected dynamics of the initial AP in Type 1 and 2 claustrum projection neurons.** A) The dynamics of the first recorded AP during step-wise current injections from Type 1 neurons (Air: 54 cells from N = 23 animals, Ethanol: 45 cells from N = 19 animals) were analyzed in air- and CIE-exposed mice. B) Phase plot of the first AP in Type 1 neurons from air- and CIE-exposed mice. C) CIE significantly lowered the threshold potential but had no effect on the AP D) peak, E) width, F) after-hyperpolarization, the G) depolarization or H) repolarization. I) The dynamics of the first recorded AP during step-wise current injections from Type 2 neurons (Air: 28 cells from N = 12 animals, Ethanol: 43 cells from N = 18 animals) were analyzed in air- and CIE-exposed mice. J) Phase plot of the first AP in Type 2 neurons from air-

and CIE-exposed mice. CIE did not affect K) the threshold potential but L) significantly reduced the peak of the AP and M) produced a trending increase in the width of the AP. CIE further N) produced a trending reduction in the after-hyperpolarization and O) a trending decrease in the depolarization rate of the AP while P) having no effect on the repolarization rate. Mean +/- SEM. Individual data points = mouse average. *p<0.05, **p<0.01, ***p<0.001, ****p<0.0001.

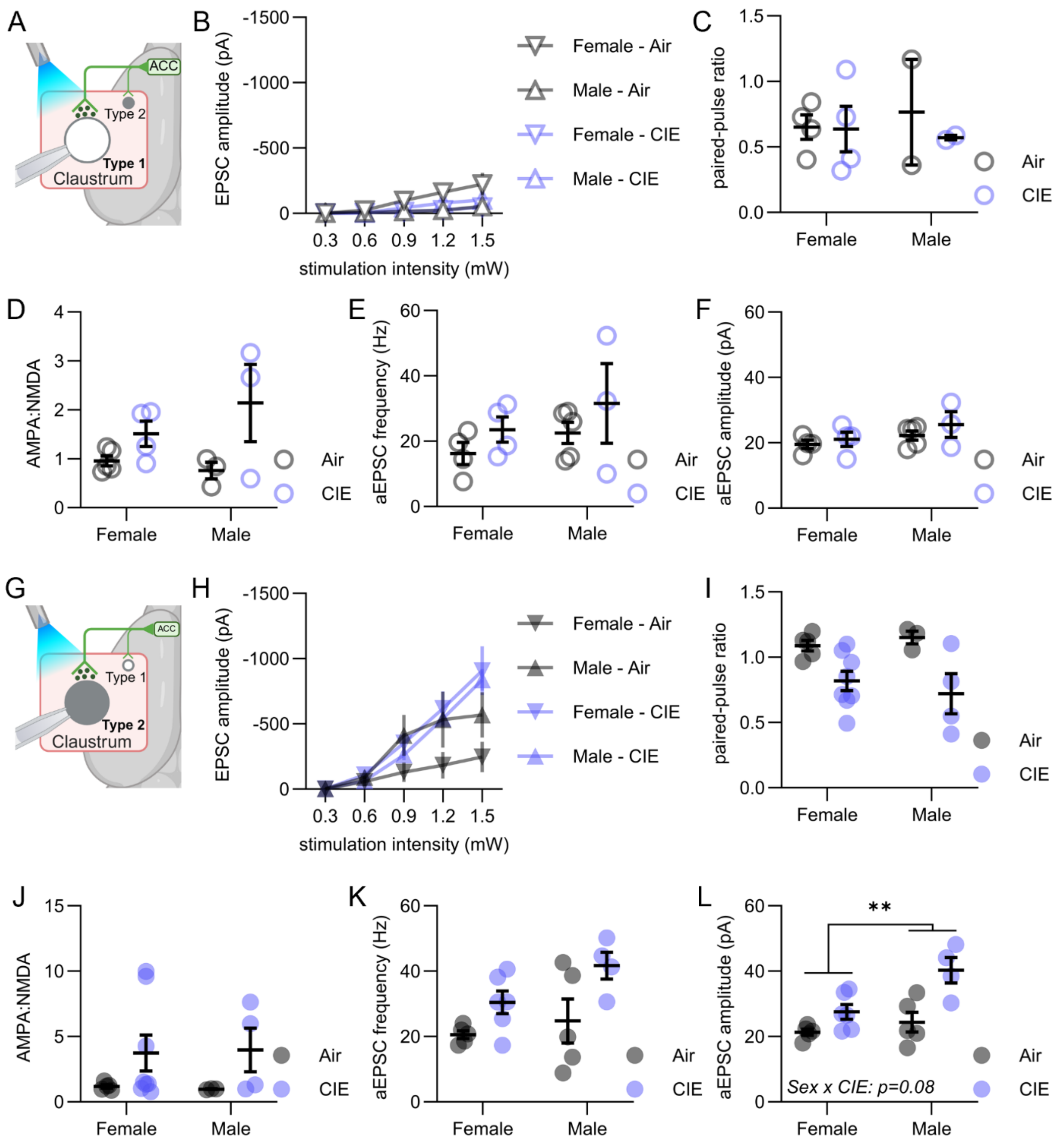


**Supplemental Figure 5: CIE sex-independently affected ACC synaptic transmission onto Type 2 claustrum projection neurons.** A) When activating ACC afferents and recording EPSCs in Type 1 claustrum neurons (Air: 14 cells from N = 5 females and 5 cells from N = 3 males, Ethanol: 13 cells from N= 4 females and 4 cells from N = 3 males), sex had no impact on B) the amplitude of the response over increasing stimulation intensities, C) the paired-pulse ratio, or C) the AMPA:NMDA ratio. When recording aEPSCs after activating ACC afferents, no effect of sex was observed on E) the frequency or F) the amplitude of aEPSCs. G) When activating ACC afferents and recording EPSCs in Type 2 claustrum neurons (Air: 16 cells from N = 6 females and

7 cells from N = 3 males, Ethanol: 19 cells from N = 8 females and 9 cells from N = 4 males), no effect of sex was observed on H) the amplitude of EPSCs over increasing stimulation intensities, I) the paired-pulse ratio or J) the AMPA:NMDA ratio. When recording aEPSCs following activation of ACC afferents, K) no effect of sex was observed on aEPSC frequency but L) a trending interaction between sex and CIE treatment was found on aEPSC amplitude and Type 2 neurons from male mice exhibited greater aEPSC amplitudes. Mean +/- SEM. Individual data points = mouse average. *$p<0.05$, **$p<0.01$, ***$p<0.001$, ****$p<0.0001$.

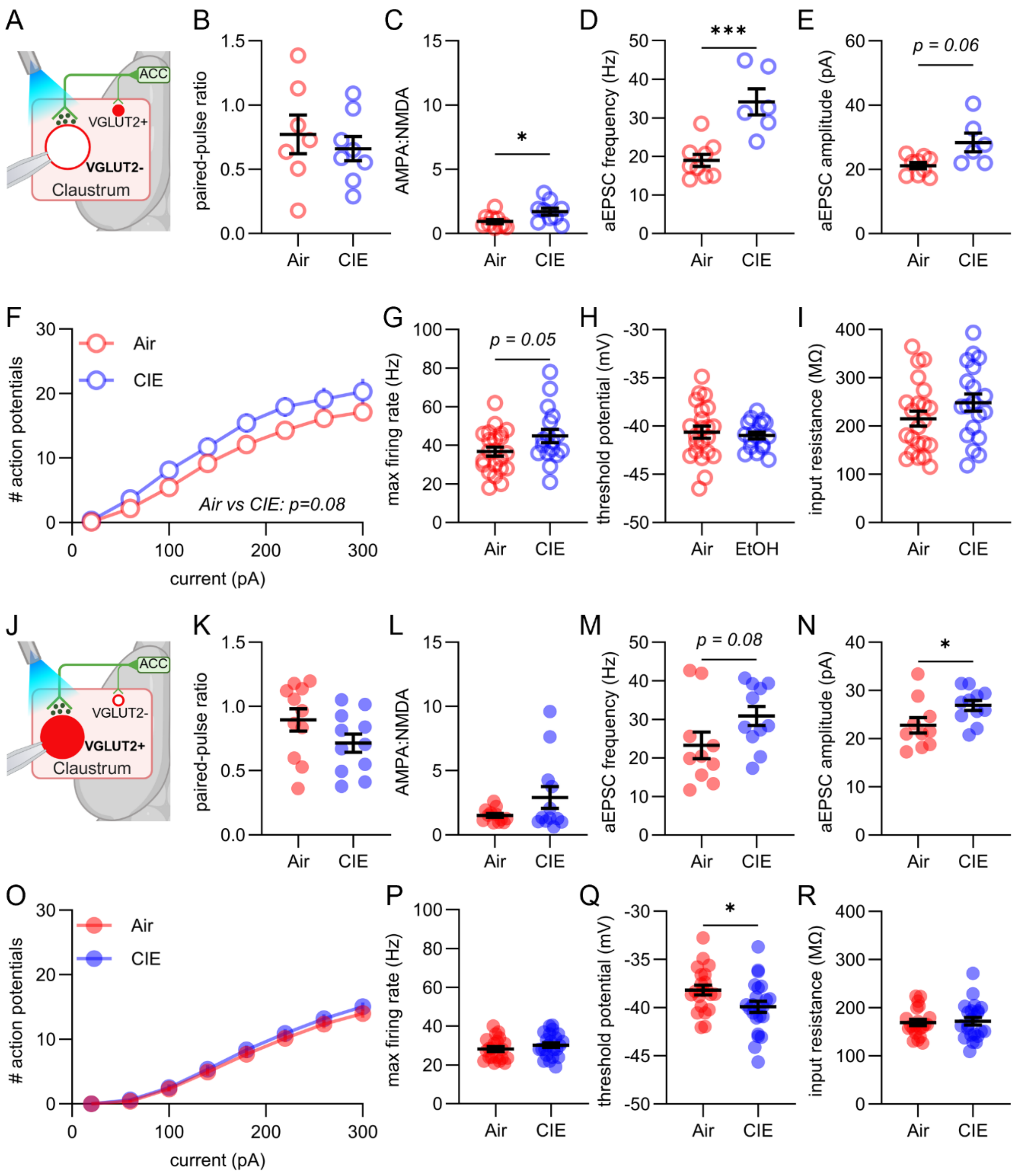


**Supplemental Figure 6: CIE-induced excitability and synaptic changes in VGLUT2- and VGLUT2+ neurons.** A) When activating ACC afferents and recording EPSCs in VGLUT2- claustrum neurons (Air: 18 cells from N = 10 animals, Ethanol: 18 cells from N = 9 animals), CIE B) did not affect the paired-pulse ratio but C) significantly increased the AMPA:NMDA ratio. When recording asynchronous EPSCs following activation of ACC afferents in VGLUT2- claustrum neurons (Air: 16 cells from N = 9 animals, Ethanol: 13 cells from N = 6 animals), CIE did not affect the D) frequency or E) amplitude of aEPSCs. F) When injecting a step-wise current into VGLUT2- neurons (Air: 39 cells from N = 22 animals, Ethanol: 36 cells from N = 17 animals), CIE produced a trending increase in AP firing with increasing current amplitude and G) resulted in a trending increase in the maximum firing rate. CIE did not affect the H) threshold potential or I) input resistance of VGLUT2- neurons. J) When activating ACC afferents and recording EPSCs in VGLUT2+ claustrum neurons (Air: 33 cells from N = 12 animals, Ethanol: 30 cells from N = 12 animals), CIE did not significantly alter the K) paired-pulse ratio or L) AMPA:NMDA ratio. When recording asynchronous EPSCs following activation of ACC afferents in VGLUT2+ neurons (Air: 27 cells from N = 10 animals, Ethanol: 29 cells from N = 11 animals), CIE M) produced a trending increase in frequency and N) significantly increased the amplitude of aEPSCs. O) When injecting a step-wise current into VGLUT2+ claustrum neurons (Air: 53 cells from N = 21 animals, Ethanol: 52 cells from N = 23 animals), CIE did not affect AP firing with increasing current amplitude or P) the maximum firing rate. Q) CIE significantly lowered the threshold potential but R) did not affect the input resistance of VGLUT2+ neurons. Mean +/- SEM. Individual data points = mouse average. *$p<0.05$, **$p<0.01$, ***$p<0.001$, ****$p<0.0001$.
